# Intra-dorsal striatal acetylcholine M1 but not dopaminergic D1 or glutamatergic NMDA receptor antagonists inhibit the consolidation of duration memory in interval timing

**DOI:** 10.1101/2021.05.24.445416

**Authors:** Masahiko Nishioka, Taisuke Kamada, Atsushi Nakata, Naoko Shiokawa, Aoi Kinoshita, Toshimichi Hata

**Author notes:** Corresponding author, 1-3 Tatara, Miyakodani, Kyotanabe, Kyoto, 610-0394, Japan.

## Abstract

The striatal beat frequency (SBF) model assumes that striatal medium spiny neurons encode duration via synaptic plasticity. Muscarinic 1 (M1) cholinergic receptors, as well as dopamine and glutamate receptors, are important for neural plasticity in the dorsal striatum. Therefore, we investigated the effect of inhibiting these receptors on the formation of duration memory. After sufficient training in a Peak interval (PI)-20 s procedure, rats were given a single or mixed infusion of a selective antagonist for the dopamine D1 receptor (SCH23390, 0.5 μg per side), the NMDA-type glutamate receptor (D-AP5, 3 μg), or the M1 receptor (pirenzepine, 10 μg) bilaterally in the dorsal striatum, immediately before starting a PI 40 s session (shift session). On the next day, the rats were tested for new duration memory (40 s) in a session in which no lever presses were reinforced (probe session). In the shift session, performance was tie, irrespective of the drug injected. However, in the probe session, the mean peak time (an index of duration memory) of the M1 + NMDA co-blockade group, but not of the D1 + NMDA co-blockade group, was lower than that of the control group (Exp. 1 and 2). In Exp. 3, the effect of the co-blockade of M1 and NMDA receptors was replicated. Moreover, sole blockade of M1 receptors induced the same effect as M1 and NMDA blockade. These results suggest that in the dorsal striatum, the M1 receptor, but not the D1 or NMDA receptors, are involved in the consolidation of duration memory.

## 1. Introduction

Animals can regulate the temporal duration of their behaviors over seconds to minutes, which is called interval timing [1]. For instance, in a fixed-interval (FI) reinforcement schedule, when a lever press is reinforced after a certain time (*t*) has elapsed from the start of the presentation of a stimulus, animals eventually learn to press the lever more frequently as *t* approaches. This behavior suggests that animals can form duration memories.

The dorsal striatum (DS) has been suggested to play an important role in duration memory. Electrophysiological studies of rats reported that the activity of the DS neurons gradually increases as the time of a reinforcement approaches [2] and different neurons in the DS fire at different time points in an interval [3]. DS lesions impair the regulation of behaviors depending on the passage of time [4]. Evidence suggests that duration memory is underpinned by neuronal plasticity in the DS. In general, memory is initially acquired as short-term memory that lasts for a few hours and then moves into long-term memory that lasts for more than a few days, through a consolidation process dependent on protein synthesis [5,6]. In fear memory, knockdown of Arc, a protein related to neural plasticity, or infusion of the protein synthesis inhibitor anisomycin, prevents memory consolidation [7,8]. In addition, in interval timing, the expression of Arc protein in the DS is changed following the formation of a new duration memory [9]. In a behavioral pharmacological study, anisomycin administered in the DS delayed the formation of new duration memories [10]. These findings suggest that plastic changes, including protein synthesis, occur in the DS at the neuronal level during the consolidation of duration memory at the behavioral level.

Synaptic plasticity in the medium spiny neurons (MSNs), which comprise the majority of neurons in the DS, occurs by sole activation or co-activation of the N-methyl-D-aspartic acid (NMDA)-type glutamate, dopamine D1, and/or muscarinic acetylcholine M1 receptors. MSNs receive glutamatergic inputs from the neocortex and dopaminergic inputs from the substantia nigra. *In vitro*, induction of LTD in MSNs by high-frequency stimulation (HFS) of cortico-striatal axons was impaired by the inhibition of D1 receptors [11]. In contrast, HFS of the axons in Mg^+^-free solution induced longterm potentiation (LTP), which was inhibited by D1 receptor blockade [12]. Moreover, activation of D1 receptors enhanced the current mediated by NMDA channels in MSNs [13]. Behavioral studies suggest that, in the DS, inhibition of NMDA receptors or extracellular signal regulated kinase (ERK), which is phosphorylated by the co-activation of D1 and NMDA receptors [14], prevents action-outcome learning in rats [15,16,17]. As to cholinergic receptors, blockade of M1 receptors suppressed the induction of LTP in the DS in Mg+-free solutions [18], and activation of M1 receptors enhanced the current mediated by NMDA receptors [19]. Taken together, results on the activation of D1, NMDA, and/or M1 receptors suggest their involvement in the plasticity of DS synapses.

Especially for the dopamine-glutamate interaction, the striatal beat frequency (SBF) model [20] theoretically supports the above idea by predicting that MSNs encode duration by the coordination of dopaminergic inputs from the substantia nigra and glutamatergic inputs from the neocortex. This model assumes that coincidental activation of glutamatergic and dopaminergic inputs to the MSNs encodes duration by changing the weight of the glutamatergic synapses. This idea is consistent with the experimental suggestion that cortico-striatal synaptic plasticity is important for the formation of duration memory and requires dopaminergic and glutamatergic inputs. Therefore, both experiment and theory suggest the importance of the role of D1 and NMDA receptors in the formation of duration memory. However, to the best of the authors’ knowledge, in the acquisition and consolidation of duration memory, the role of not only the interaction of NMDA receptors with D1 in the DS, but also that of M1 receptors has not been investigated.

To investigate the role of D1 and M1 receptor interactions with NMDA receptors in the DS in the acquisition and consolidation of duration memory, we applied the “time-shift paradigm” [21] with a probe session [22] to the peak interval (PI) procedure. The PI procedure consists of food and empty trials presented in a random order and separated by an inter-trial interval (ITI). Each trial starts with the presentation of a signal stimulus, such as a light or tone. In a food trial, the first lever press after a required time (e.g., 20 s) is reinforced, and then the trial ends. In an empty trial, no lever press is reinforced and the trial ends after a predetermined time (e.g., 60 s). In empty trials, when the number of lever-press responses was plotted as a function of time elapsed from trial start, a bell-shaped distribution of response rate was observed, with the maximum number of responses occurring around the required time. After acquisition under a first required time, the required time is then abruptly changed (e.g., to 40 s) in a “shift session” In the shift session, the response rate distribution shifts toward the new required time, suggesting the formation of a new duration memory (time shift paradigm). It has been shown that the infusion of protein synthesis inhibitors into the DS delayed the acquisition of new duration memory [11], and that Arc expression changed following the formation of a new duration memory [10]. These previous studies suggest that the time-shift paradigm is suitable for examining the acquisition of duration memory. To study the consolidation of duration memory, we added a “probe session” following the shift session, in which all trials were empty trials and drugs were not administered [22]. Impairment of consolidation is operationally defined as the appearance of impairment in long-term memory with no impairment in short-term memory when the drug is infused before or after the session [6]. Recently, it has been reported that suppression of dorsal hippocampal activity by muscimol did not affect the acquisition of new duration memory in the shift session but impaired the memory if tested in a probe session the next day without the drug [22]. These findings suggest that short-term memory from recent food trials guided adaptive behavior in the shift session, while long-term memory, which should have been observed in the probe session, did not form. This interpretation is consistent with the operational definition of impaired memory consolidation. Therefore, we believe that the present procedure can be used to examine the effects of drugs specifically on the consolidation of duration memory.

The present study aimed to investigate the effect of single or co-infusions of D1, NMDA, and/or M1 receptor inhibitors into the DS on the acquisition and consolidation of duration memory using a time-shift paradigm with a probe session with a peak-interval task. If the peak of the response rate distribution shifts toward the new required duration in the shift session but returns to the baseline level in the probe session, the suggestion would be that the drug(s) prevent(s) consolidation but not acquisition of duration memory. Our findings suggest that sole infusion of an M1 receptor blocker inhibits the consolidation of duration memory.

## 2. Experiment 1: Effect of the intra-DS mixed infusion of D1 and NMDA receptor antagonists on the acquisition and consolidation of duration memory

The SBF model assumes that the interaction between dopamine and glutamate receptors in the DS is important for the formation of duration memory. In particular, D1 dopamine and NMDA type glutamate receptors are known to be important for neuronal plasticity in the DS. Therefore, in Experiment 1, we examined the effect of intra-DS infusion of antagonists for these receptors on the acquisition and consolidation of duration memory. All experimental procedures were approved by the Doshisha Committee of Animal Experiments (A18077).

### 2.1. Materials and Methods

#### 2.1.1. Subjects

Twenty experimentally naïve, male Wistar albino rats, aged approximately 11 weeks, were used. They were housed individually in cages. To enhance their drive for food, their body weights were decreased to 85% of their free-feeding weights by the start of the experiment. Considering natural growth, body weight was allowed to increase by 5 g each week. The rats were allowed free access to water in their cages. A 12:12 light-dark cycle was used, and the light phase started at 8:00 AM. The experiment was performed during the light phase.

#### 2.1.2. Apparatus

Eight identical operant chambers were individually enclosed in a sound-proof box equipped with a 0.8-W LED and a buzzer (M2BJ-B24, OMRON, Kyoto, Japan) on the ceiling. A non-retractable lever and a food cup were attached to the left and the center of the front panel (40 mm and 15 mm above the floor grids, respectively). The floor grids were made with stainless steel bars (3 mm in diameter) separated by 10 mm. A custom application developed using XOJO ^®^ (XOJO Inc. Austin, TX, USA) running on two PowerBook Air computers (Apple, Cupertino, CA, USA) controlled the experiment and collected the data. They were interfaced with two programmable controllers (SYSMAC CPM1A40CDR-A-V1, OMRON, Kyoto, Japan) and two USB I/O controllers (RBIO-2U, Kyoritsu Electronic, Osaka, Japan).

#### 2.1.3. Procedure

##### 2.1.3.1. Shaping

After handling for 5 min per day for five days, the rats were habituated to the operant chamber for 10 min. They were then trained to press the lever to earn food pellets (F0021-J, Flemington, NJ, USA) under a continuous reinforcement schedule until they had pressed the lever 60 times, or 20 min passed in each session. The light of the sound-proof box was turned on during the session. The acquisition criterion was that they pressed the lever 60 times within 20 min in three successive sessions.

##### 2.1.3.2. Training (PI-20 s, sessions 1–30)

After completion of the shaping, they were trained in the PI-20 s procedure with one session per day. Each session consisted of food and empty trials. Both trials started with the onset of the light and buzzer and finished with their offsets. In the food trials, the first lever press after a required time (e.g., 20 s) was reinforced, and then the trial ended. (discrete trial FI-20 s). In an empty trial, no lever press was reinforced, and the trial ended after a predetermined time (e.g., 60 s). After finishing each trial, the intertrial interval (ITI) for 40 ± 10 s began. If the lever was pressed within the last 5 s of the ITI, the ITI was extended for 10 s from the lever press. Two types of trials were presented in pseudo-random order so that the empty trials were not presented in four consecutive trials. The ratio of the food and empty trials was 42:18 until session 25 and 15:15 after session 26.

##### 2.1.3.3. Surgery

The rats were anesthetized with 2%–3% isoflurane (2–3 L/min flow rate, MK-A110, Muromachi, Tokyo, Japan) by inhalation. They were then fixed on a stereotactic flame (David Kopf Instruments, Tujunga, CA, USA). Based on a brain map [23], they were bilaterally implanted with a guide cannula (C232G-5.0/SPC, Plastic One, Roanoke, VA, USA) into the DS (from bregma, AP: +0.5 mm, ML: ±2.5 mm, DV: −3.8 mm from skull). The cannula was fixed to the skull with dental cement and two small screws. The rats were then inserted with a dummy cannula (C232DC-5.0/SPC, Plastic One) into the guide cannula and fixed with a dust cap (363DC, Plastic One). To avoid infection, an antibiotic agent (Mycillinsol, Meiji, Tokyo, Japan) was applied to the surgical sites once a day for three days after surgery. After a week of recovery period after surgery, the rats were trained in re-training sessions.

##### 2.1.3.4. Re-training (PI-20 s, sessions 31–36)

After the recovery period, the rats were re-trained using a procedure similar to the training sessions (26–30) for six sessions. The last session of the training sessions was defined as the baseline session.

##### 2.1.3.5. Shift session (PI-40 s, sessions 37)

On the day after the baseline session, each rat was assigned to one of two groups, the artificial cerebrospinal fluid (aCSF) or drug groups, so that the performances of the six re-training sessions became even. Immediately before starting each session, the dummy cannula was replaced with an injection cannula (C2321-5.0/SPC, Plastic One), which extended 1 mm from the tip of the guide cannula for infusion. ACSF or SCH23390 (D1 receptor selective antagonist, 0.5 μg/side) and D-AP5 (NMDA type glutamate receptor selective antagonist, 3.0 μg/side) were infused into the bilateral DS for each group using a gastight syringe (100 μL, Hamilton, Reno, NV, USA) and a micro syringe pump (ESP-32, Eicom, Kyoto, Japan). The injection lasted for two min at a flow rate of 0.5 μL/min. The injection cannula was left in place for one min to allow dispersion of the solution. Then, the injection cannula was replaced with the dummy cannula. Importantly, PI-40 s was executed in the shift session, in which a lever press 40 s after the start of the trial was reinforced in a food trial. The empty trial lasted 120 seconds. The food and empty trials were respectively presented 15 times. The other parameters were the same as those used in the training sessions.

##### 2.1.3.6. Probe session (sessions 38)

On the day after the shift session, the rats were tested in a probe session without infusion. In the probe session, only 15 empty trials, each of which was the same as the empty trial of the shift session, were included. The other parameters were the same as those used in the shift session.

#### 2.1.4. Dependent variables

##### 2.1.4.1. Session-by-session analysis

Details of the data analysis were performed according to a previous study [22]. Data from empty trials were used for the analysis. Briefly, as an index of the subjective length of the target duration, we calculated the peak time. The total number of leverpress responses in the 3-s bin (such as 0–3 s, 1–4 s) in each session were counted individually. The total number of responses in each bin was individually converted to a percentage of the maximum number of responses (response rates). The response rate distributions were fitted to the Gaussian curve, *R(t)* = *a + b exp{-(t-c)^2^/d^2^*} with a fitting function (MATLAB, ver. R. 2020a, MathWorks, MA, USA). *R(t)* is the estimated response rate for each bin. *c* is a putative parameter of bin having the maximum response rate and is thus defined as the peak time, which is an index of the subjective length for the target duration. The initial values of *a, b*, and *d* were set to 0, 100, and 8, respectively, for all subjects. To determine the initial *c*, the first and last bins, which had response rates of over 85%, were detected. The mean of the two class values of these bins was set as the initial c. To fit the curve, the data in the range of 1.5 s to 38.5 s was used in the re-training session. In the shift and the probe sessions, the range was 1.5 s to 78.5 s. Curve fitting was also performed for the group means of the response rate distributions. All bin data were used for the fitting of the group mean.

As an index of the precision for the subjective length of the target duration, the discrimination index (*DI*) was calculated using *the maximum response rate 100(%)/mean response rate over all bins*.

As an index of the frequency of lever-press behavior during the session, the number of responses per second was calculated as *the total number of responses/the total sec of all empty trials*.

##### 2.1.4.2. Trial-by-trial analysis

For each empty trial, the start and stop times were calculated. In an individual empty trial, rats typically started pressing the lever before the required time, kept responding, and then stopped responding after the required time. The time points of the transition from low to high and high to low can be, respectively, detected as the start time and stop time by the algorithm proposed in a previous study [24]. The spread was calculated as the stop time minus the start time. A trial with only a response was excluded from the data analysis.

#### 2.1.5. Histology

After completing the probe session, rats were infused with dye (2% pontamine sky blue) via injection cannula and perfused intracardially with saline and ALTFiX^®^ (FALMA, Tokyo, Japan) under deep anesthesia with sodium pentobarbital (130 mg/kg, i. p.). Their brains were removed and soaked in 10% and 30 % sucrose phosphate buffer (0.1M) until they sunk in each solution. The brains were sectioned coronally using a cryostat (CM1850, Leica, Wetzlar, Germany) to 40 μm, mounted on glass slides, and stained with cresyl violet.

#### 2.1.6. Statistical analysis

We used two-way mixed ANOVAs followed by the test of simple main effects and Bonferroni’s multiple comparisons with the anovakun function (ver. 4.8.5., http://riseki.php.xdomain.jp/index.php) in R software (ver. 4.0.3, R Core Team, Vienna, Austria) [25]. The adjusted *p*-values in the multiple comparisons were manually calculated by the authors based on the analysis log. The significance level of the statistical analyses was set at *α* = .050.

### 2.2. Results

Two rats were removed from the analysis: one whose *R^2^* in the probe session was below −2 *SD* and the other who failed to form an appropriate peak curve for target duration throughout the PI-20 s sessions. As a result, the number of subjects was eight in the aCSF and 10 in the SCH23390+D-AP5 group (SCH+AP5 group).

#### 2.2.1. Histology

The locations of the internal cannula tips are shown in Figure 1B. All locations were within the DS. The brain sections of two rats in the aCSF group were missed; hence, we evaluated their guide cannulas and referred to the surgical records. There was no damage in the cannulas, and the recorded coordinates were consistent with those of the other animals. Therefore, the two rats were included in the behavioral analysis.

**Figure 1.**
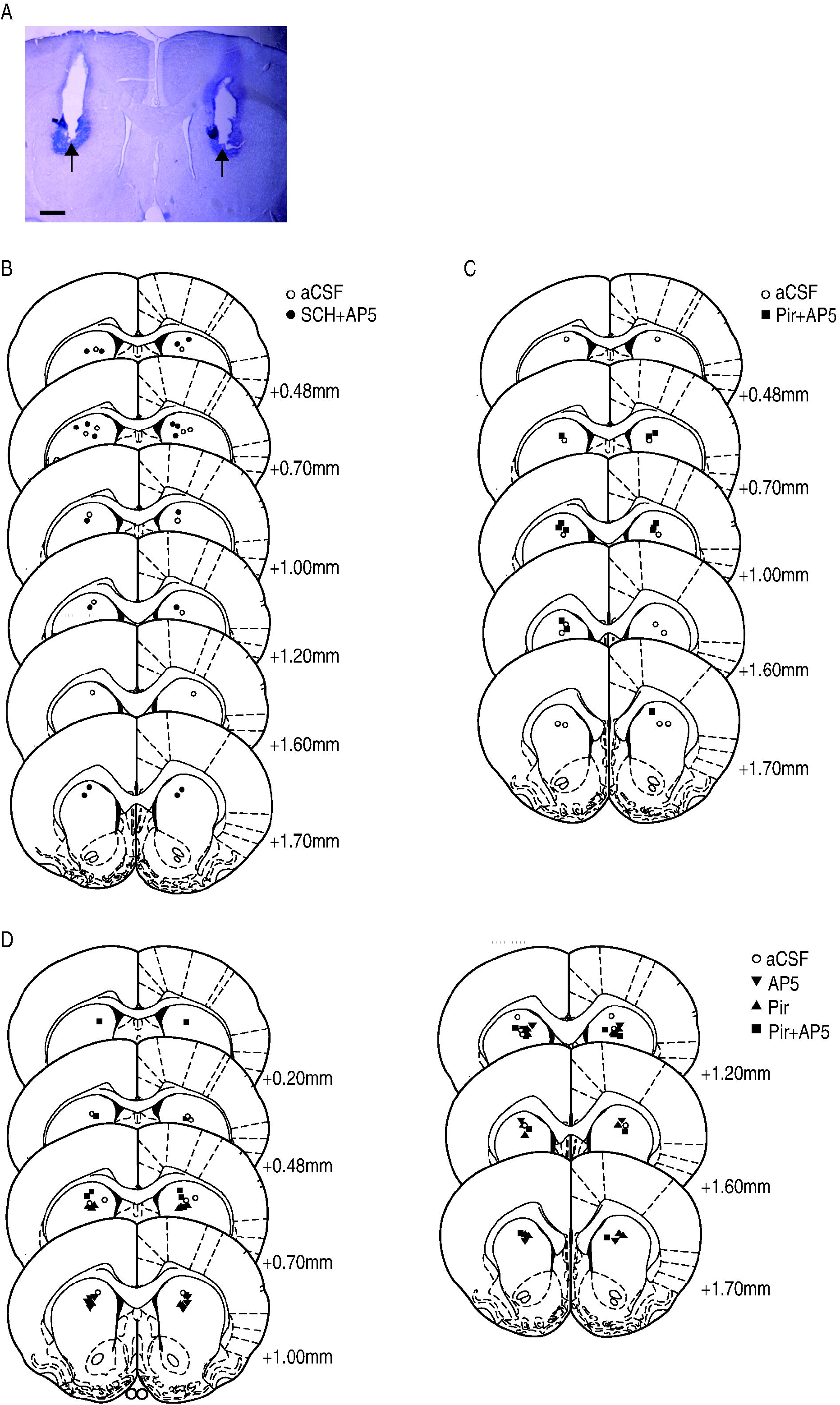
A representative image and the position of cannula. **(A)** A representative image showing the position of cannula insertion. Black arrows indicate tips of the cannula. Scale bar = 1 mm. **(B-D)** The locations of the cannula tips in Experiment 1 **(B)**, 2 **(C)**, and 3 **(D)**. Open circles denote the aCSF group and closed squares the Pir+AP5 group. Closed circles denote the SCH+AP5 group, closed inverted triangles the AP5 group, and closed triangles the Pir group. Reprinted from Paxinos, G. & Watson, C. *The Rat Brain in Stereotaxic Coordinates. 4th ed. [CD-ROM]*, Copyright (1998) [23].

#### 2.2.2. Session by session analysis

##### 2.2.2.1. Response rate distributions

The mean response rate distributions and their fitted curves are shown in Figure 2. In the baseline session (Figure 2A), the distributions of both groups overlapped with peaks at approximately 18 s. In the shift session (Figure 2B), the distributions shifted rightward compared to the baseline session. The peaks in both groups were located at approximately 28 s. In the probe session (Figure 2C), the distribution was in the middle of 20 and 40 s. The peaks in both groups were located at approximately 25 s. The mean *R^2^s* (± *SEM*) values of the fitting curve are shown in Table 1. The values were greater than .818 in all sessions. A mixed two-way ANOVA (group × session) showed a significant main effect of session [*F* (2,32) = 11.192, *p* < .001, *ηG^2^*= .306], but not the main effect of the group and the interaction [*F* (1,16) = 1.795, *p* = .199, *ηG^2^*= .040; *F* (2,32) =.395, *p* = .677, *ηG^2^* = .015, respectively]. In multiple comparisons for the session, the *R^2^s* of the shift and probe sessions was significantly lower than that of the baseline session [*p_s_* < .001].

**Figure 2.**
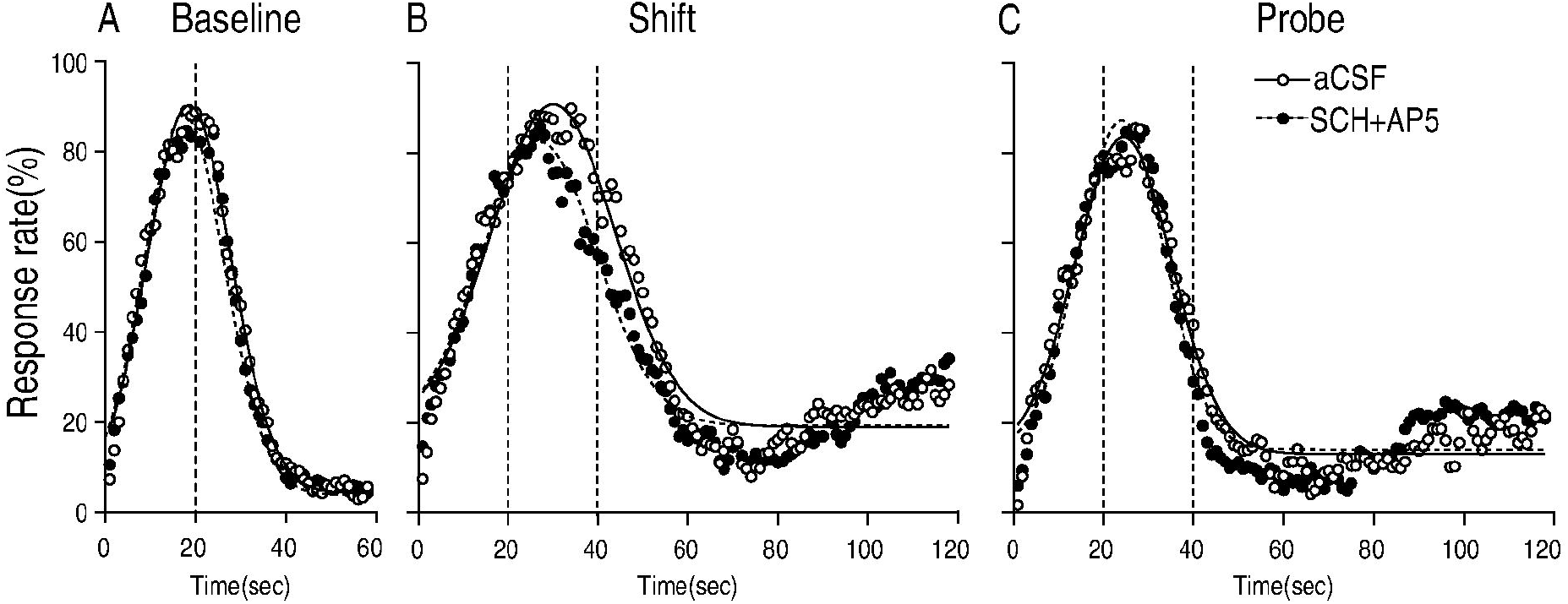
Mean response rate distributions and fitted Gaussian curves. Shown are baseline **(A)**, shift **(B)**, and probe sessions **(C)**. Open circles denote the aCSF group, and closed circles the SCH+AP5 group. The vertical dotted lines show the required (reinforced) times of the PI-20 s and PI-40 s sessions.

**Table 1.**
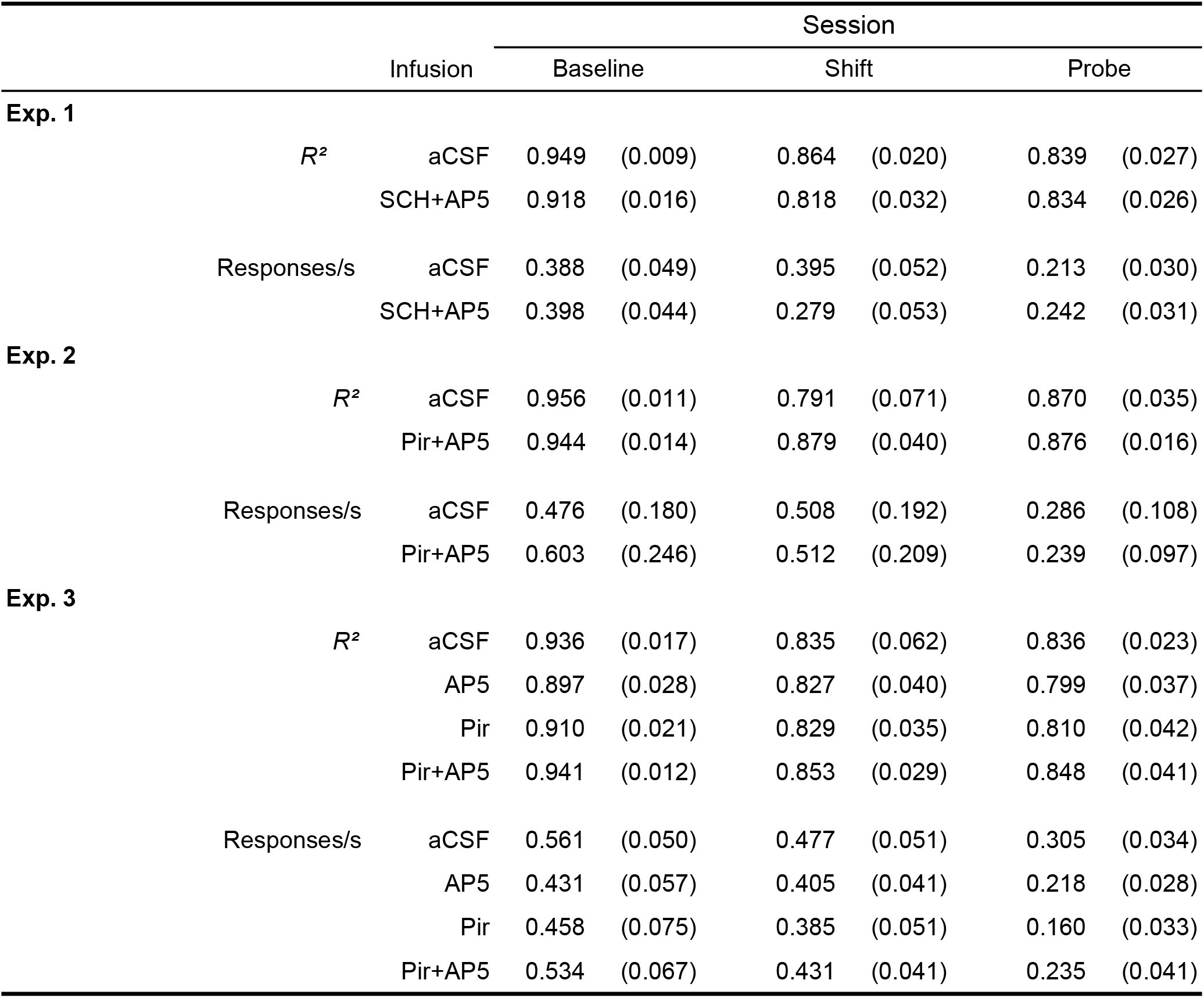
Mean (±*SEM*) of *R^2^* and the number of responses per second.

##### 2.2.2.2. Peak time

The mean peak times (± *SEM*) from the first re-training session to the probe session are shown in Figure 3A. During the training sessions, the mean peak times of both groups were consistently approximately 18 s. A mixed two-way ANOVA in the retraining sessions showed no significant main effect of the group [*F* (1,16) = .366, *p* = .554, *ηG^2^* = .013], the session [*F* (2.91, 46.54) = 1.897, adjusted by Greenhouse-Geisser’s epsilon, *p* = .145, *ηG^2^* = .048], and interaction [*F* (2.91, 46.54) = 1.479, *p* = .233, *ηG^2^* = .037]. The mean peak times of both groups in the shift and probe sessions were shifted in the same way. A mixed two-way ANOVA from the baseline to the probe sessions revealed a significant main effect of session [*F* (2,32) = 35.131, *p* < .001, *ηG^2^* = .601]. In multiple comparisons, all combinations showed significant differences [*p_s_* < .047]. The main effect of the group and interaction was not significant [*F* (1,16) = 2.089, *p* = .168, *ηG^2^* = .039; *F* (2,32) = .562, *p* = .576, *ηG^2^* = .024, respectively].

**Figure 3.**
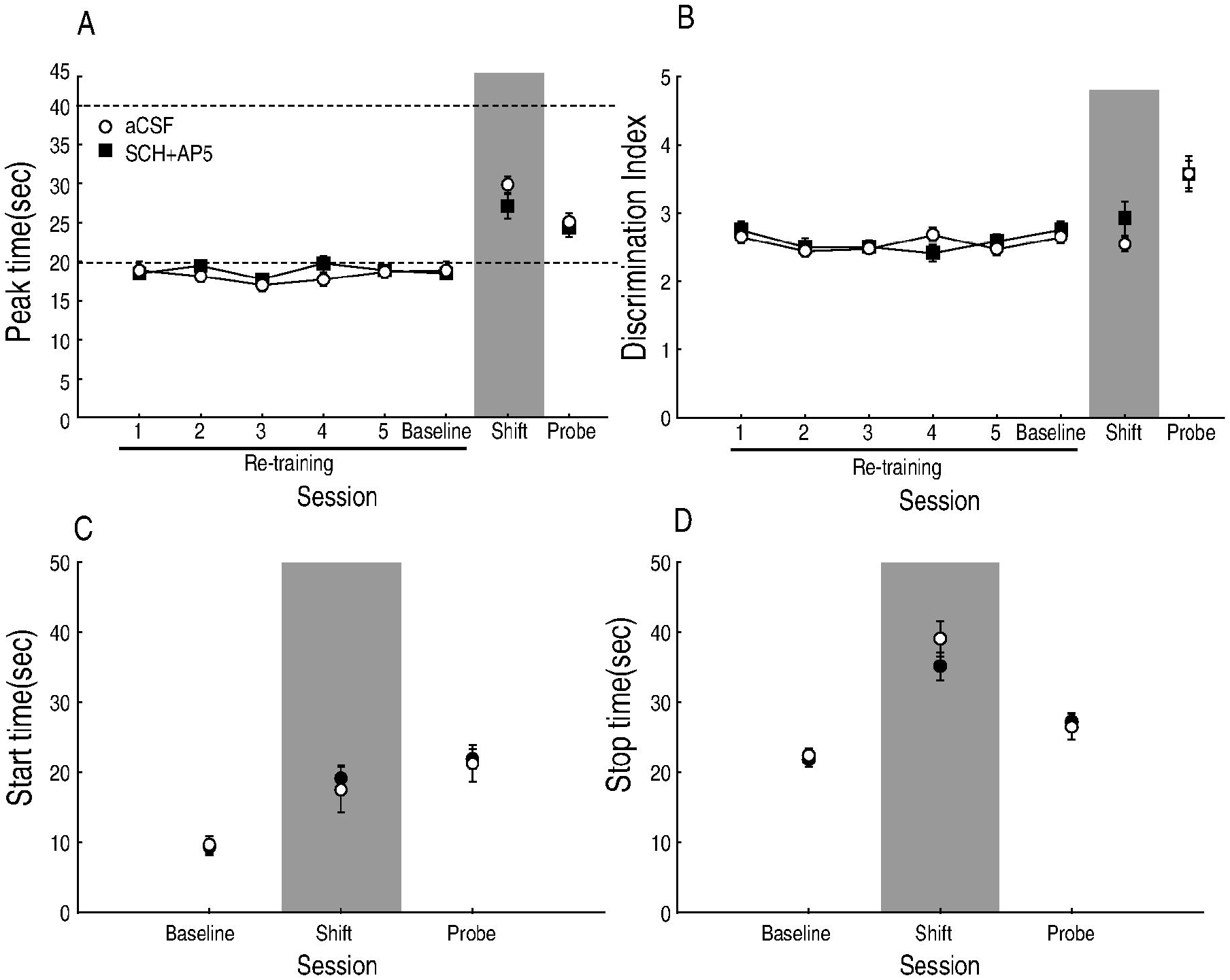
Mean peak time, discrimination index (*DI*), start, and stop time (±*SEM*). **(A)** Mean peak time (*±SEM*) from the re-training to the probe session. Open circles denote the aCSF group, and closed circles the SCH+AP5 group. **(B)** Mean *DI* ± *SEM* from the retraining to the probe session. Mean (±*SEM*) start **(C)** and stop **(D)** times at the baseline, shift, and probe sessions.

##### 2.2.2.3. DI

The mean *DI*s (± *SEM*) from the first re-training session to the probe session are shown in Figure 3B. Although the mean *DI*s varied slightly, there was no difference between the groups. A mixed two-way ANOVA in the re-training sessions showed a significant main effect of session [*F* (2.26,36.11) = 3.302, *p* = .043, *ηG^2^* = .083], but the main effect of the group and the interaction were significant [*F* (1,16) = .030, *p* =.864, *ηG^2^* = .001; *F* (2.26,36.11) = 1.676, *p* = .199, *ηG^2^* = .044, respectively]. There was no noticeable difference between the groups in the mean *DI*s in the baseline, shift, and probe sessions. A mixed two-way ANOVA from the baseline to the probe sessions revealed a significant main effect of session [*F* (2,32) = 19.955, *p* < .001, *ηG^2^* = .360], but not the main effect of the group and the interaction [*F* (1,16) = .627, *p* = .440, *ηG^2^* = .021; *F* (2,32) = .809, *p* = .454, *ηG^2^* = .022, respectively]. In multiple comparisons of the session, the *DI* of the probe session was significantly higher than that of the baseline and shift sessions [*p_s_* < .001].

##### 2.2.2.4. The number of responses per sec

The mean number of responses per second (± *SEM*) in the baseline, shift, and probe sessions are shown in Table 1. There were no noticeable differences between the groups in the baseline and probe sessions. A mixed two-way ANOVA revealed a significant main effect of the session and the interaction [*F* (2,32) = 17.779, *p* < .001, *ηG^2^* = .227; *F* (2,32) = 3.864, *p* = .031, *ηG^2^* = .060, respectively], but not the main effect of the group [*F* (1,16) = .226, *p* = .641, *ηG^2^* = .010]. Significant simple main effects of the session were detected in both groups [*F* (2,14) = 8.365, *p* = .004, *ηG^2^* = .334; *F* (2,18) = 14.118, *p* < .001, *ηG^2^* = .204, respectively]. In multiple comparisons, the value of the probe session was significantly lower than that of the shift session in the aCSF group [*p* = .040]. In the SCH+AP5 group, the values of the shift and probe sessions were significantly lower than those in the baseline session [*p_s_* < .010].

#### 2.2.3. Trial by trial analysis

The mean start and stop times (± *SEM)* are shown in Figure 3C and D, respectively. In both indices, there were no noticeable differences between the groups in the baseline, shift, and probe sessions. A mixed two-way ANOVA of the start time showed a significant main effect of the session [*F* (2,32) = 22.772, *p* < .001, *ηG^2^* = .460], but not the main effect of the group and the interaction [*F* (1,16) = .121, *p* = .732, *ηG^2^* = .003; *F* (2,32) = .137, *p* = .873, *ηG^2^* = .005, respectively]. In multiple comparisons of the sessions, the start time of the shift and probe sessions was significantly higher than that of the baseline session [*p_s_* < .001]. For the stop time, the main effect of the session was significant [*F* (2,32) = 61.688. *p* < .001, *ηG^2^* = .637], but not the main effect of the group and the interaction [*F* (1,16) = .554, *p* = .467, *ηG^2^* = .019; *F* (2,32) = 1.546, *p* = .229, *ηG^2^* = .042, respectively]. In multiple comparisons of sessions, all combinations showed significant differences [*p_s_* < .006].

### 2.3. Discussions

The results of the shift session suggest that the co-inhibition of the D1 and NMDA receptors did not impair the acquisition of new-duration memory. The peak times, start time, and stop time of both groups significantly increased from baseline to the shift session (Figure 3A, C, and D). The peak time was located at approximately in the middle of 20 s and 40 s. A behavioral study reported that when rats started to learn a new required time after learning an original time in the PI procedure, the peak time was located at the middle of the old and new required time during several dozen trials [26]. Therefore, the peak times in the shift session suggest that rats in both groups learned the new required time during the shift session. Moreover, there were no significant differences between the groups in these indices. These results suggest that the 1) new duration memory was acquired in both groups, and 2) the strength of acquired memory was similar to each other. The between-group indifference of the *DIs* suggests that the precision of the subjective length of the duration was also not affected by the drugs (Figure 3B).

More importantly, the results of the probe session suggest that the co-inhibition of D1 and NMDA receptors did not impair the consolidation of new-duration memory. In the probe session, there was no significant difference in the peak time, start time, and stop time (Figures 3A, C, and D). Namely, the infusion of drugs before the shift session did not impair the long-term memory tested in the probe session or the short-term memory tested in the shift session. In contrast, the impairment of consolidation is operationally defined as the appearance of impairment in long-term memory, with no impairment in short-term memory, when the drug is infused before or after the acquisition session [6]. Our findings were inconsistent with this definition. Moreover, there was no difference in the DIs between the groups (Figure 3B). This means that the infusion of drugs before the shift session did not impair the precision of the subjective length of the duration in the probe session.

The low concentration of the drugs cannot explain the lack of impairment in memory formation. Our doses were 0.5 μg and 3 μg/side for SCH23390 and D-AP5, respectively. The number of lever presses per second of the aCSF did not change from the baseline to the shift session, whereas that of the SCH+AP5 group was significantly decreased (Table 1). This finding suggests that the drugs affected the lever-press behavior. Previous studies using other tasks also suggest that the infusion of the same or lower concentrations of these drugs impaired several kinds of learning or behaviors; the intra-accumbens core co-infusion of SCH23390 (0.09 μg) and D-AP5 (0.1 μg) impaired lever-press learning [27]. For the sole infusion of D-AP-5, an intra-ventral striatum infusion of D-AP5 (3 μg) impaired left-right discrimination learning, whereas an intra-dorsal striatum infusion of the same dose of the drug extended the latency without affecting learning [28]. An intra-DS D-AP5 impaired the acquisition of spatial learning in an eight-arm radial maze (1 μg) [29], consolidation in a cued water maze task (2 μg) [30], and action-outcome learning (0.5 μg) [15]. For the sole infusion of SCH23390, an intra-DS SCH23390 infusion reduced the pre-pulse inhibition (0.4 μg) [31] and impulsive behavior (0.1 μg) [32]. These findings suggest that the doses used in our study were sufficient to modulate neural activity and behavior. Moreover, there was no evidence of impairment of memory formation in our results. Therefore, it is suggested that the co-inhibition of D1 and NMDA receptors in the DS does not impair the formation of memory.

## 3. Experiment 2: Effect of the intra-DS mixed infusion of M1 and NMDA receptor antagonist on the acquisition and consolidation of duration memory

In addition to D1 receptors, muscarinic M1 acetylcholine receptors interact with NMDA receptors in DS neuronal plasticity. In Experiment 2, the effect of the intra-DS mixed infusion of M1 and NMDA receptor antagonists on the acquisition and consolidation of duration memory was examined. All experimental procedures were approved by the Doshisha Committee of Animal Experiments (A17075).

### 3.1. Materials and Methods

The materials and methods were the same as in Experiment 1, except for the following points. Another cohort of 16 naïve male Wistar albino rats, aged approximately 11 weeks, was used. In training, the ratio of the food and empty trials was 42:18 until session 21 and 15:15 after session 22. In the shift session, aCSF or pirenzepine, an M1 receptor selective antagonist, (10 μg/side) and D-AP5 (3.0 μg/side) were infused.

### 3.2. Results

Three rats were removed from the analysis: one whose *R^2^* in the probe session was below −2 SD, one that failed an appropriate peak time estimation for the target duration in the PI-20 s sessions, and one that failed to form bell-shaped curves for target duration throughout the PI-20 s sessions. As a result, there were seven subjects in the aCSF and six in the pirenzepine+D-AP5 (Pir+AP5) group.

#### 3.2.1. Histology

All internal cannula tips are shown in Figure 1C. All internal cannulas were inserted into the DS.

#### 3.2.2. Session by session analysis

##### 3.2.2.1. Response rate distributions

The mean response rate distributions and their fitted curves for both groups are shown in Figure 4. In the baseline session (Figure 4A), the distributions of both groups overlapped with peaks at approximately 19 s. In the shift session (Figure 4B), the distributions shifted rightward compared to the baseline session. The peaks in both groups were located at approximately 32 s. In the probe session (Figure 4C), the distribution of the Pir+AP5 group was located leftward compared to that in the aCSF group. The peaks of the Pir+AP5 group were located at approximately 22 s, whereas those of the aCSF group were located at approximately 27 s. The mean *R^2^s* (± *SEM*) values of the fitting curve are shown in Table1. The values were over .791 in all the sessions. A mixed two-way ANOVA showed a main effect of session [*F* (2,22) = 5.907, *p* = .009, *ηG^2^* = .213], but not the main effect of the group and the interaction [*F* (1,11) = .484, *p* = .501, *ηG^2^* = .021; *F* (2,22) = 1.219, *p* = .315, *ηG^2^* = .053]. In multiple comparisons, the *R^2^* of the probe session was lower than that of the baseline session [*p* = .008].

**Figure 4.**
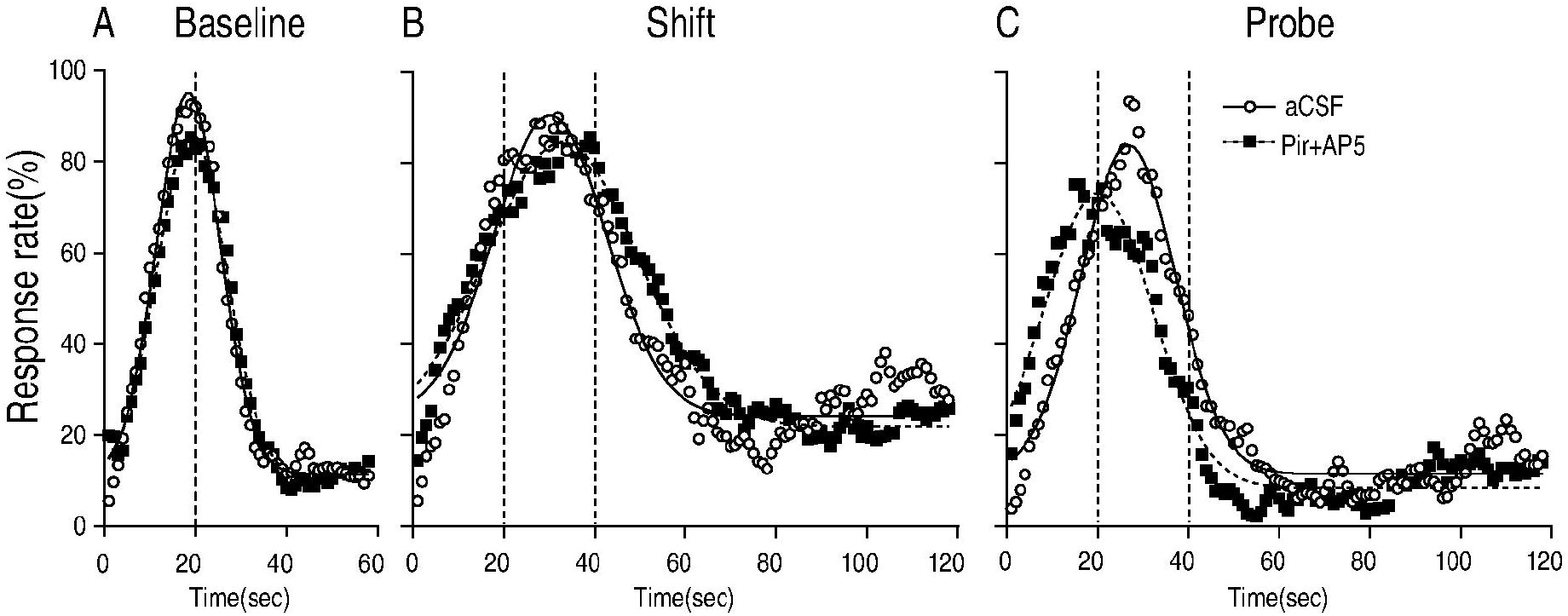
Mean response rate distributions and fitted Gaussian curves. Shown are baseline **(A)**, shift **(B)**, and probe sessions **(C)**. Open circles denote the aCSF group, and closed squares denote the Pir+AP5 group. The vertical dotted lines show the target durations of the PI-20 s and PI-40 s sessions.

##### 3.2.2.2. Peak time

The mean peak times (± *SEM*) from the first re-training session to the probe session are shown in Figure 5A. During the training sessions, the mean peak times of both groups were consistently approximately 18 s. A mixed two-way ANOVA of the peak times in the retraining sessions showed a significant main effect of session [*F* (2.38,26.23) = 9.311, *p* < .001, *ηG^2^* = .202], but not the main effect of the group and the interaction [*F* (1,11) = .002, *p* = .970, *ηG^2^* < .001; *F* (2.38,26.23) = .483, *p* = .655, *ηG^2^* = .013, respectively]. The mean peak times in the baseline and shift sessions were not different between the groups, whereas that of the Pir+AP5 group in the probe session was lower than that in the aCSF group. A mixed two-way ANOVA from the baseline to the probe sessions revealed a significant main effect of the session and the interaction [*F* (2,22) = 39.056, *p* < .001, *ηG^2^* = .585, *F* (2,22) = 5.342, *p* = .013, *ηG^2^* = .162, respectively], but not the main effect of group [*F* (1,11) = .224, *p* = .645, *ηG^2^* = .012]. Importantly, the simple main effect of the group was significant at the probe session [*F* (1,11) = 5.786, *p* = .035, *ηG^2^* = .345]. The simple main effects of the session were also significant for the aCSF and Pir+AP5 groups [*F* (2,12) = 24.142, *p* < .001, *ηG^2^* = .680; *F* (2,10) = 19.748, *p* < .001, *ηG^2^* = .573]. In multiple comparisons, the peak time of the shift and probe sessions was significantly higher than that of the baseline session in the aCSF group [*p_s_* < .009]. In the Pir+AP5 group, the peak time of the shift session was significantly higher than that of the baseline and probe sessions [*p_s_* < .038]. Importantly, the difference between the baseline and probe sessions was not significant.

**Figure 5.**
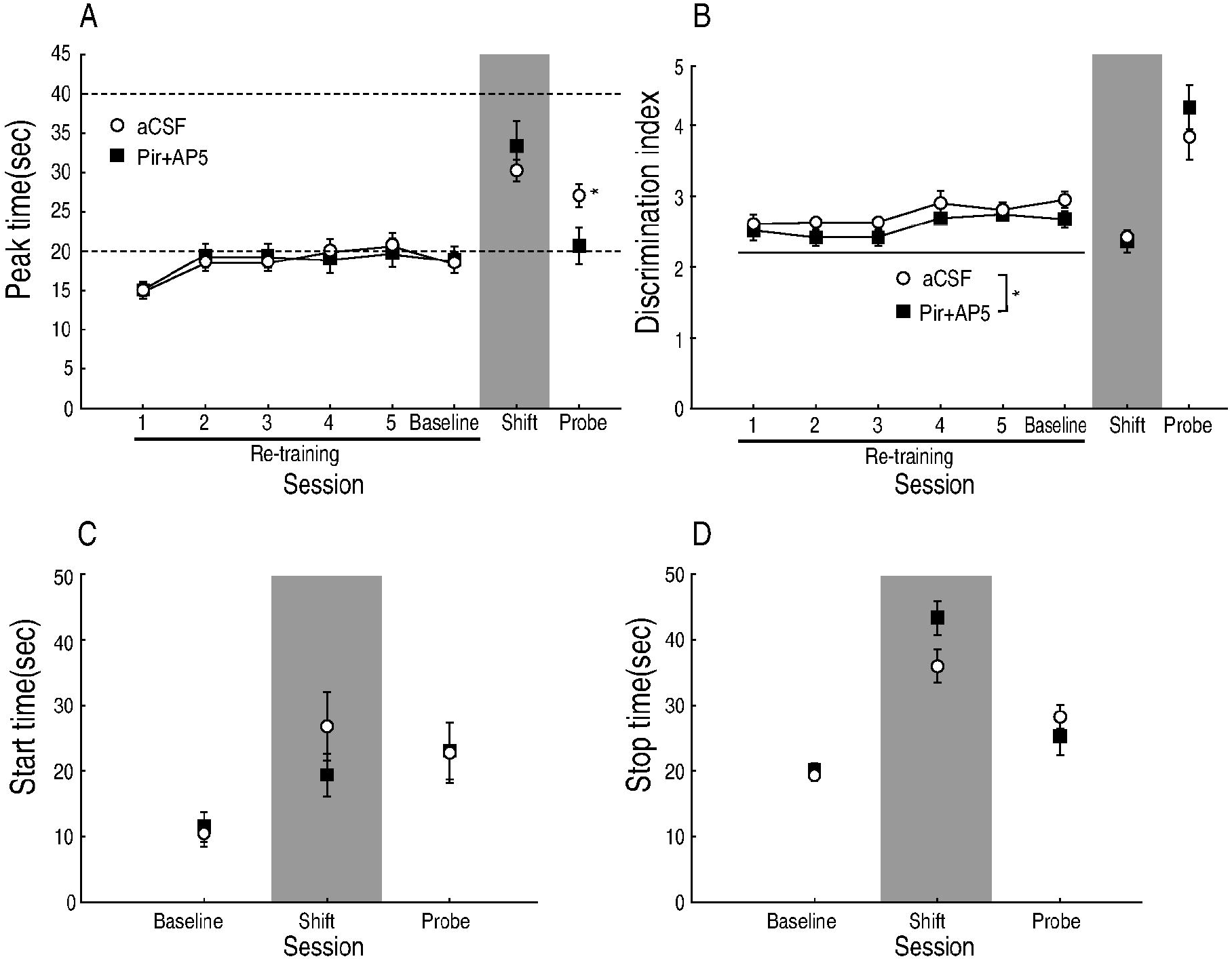
Mean peak time, discrimination index (*DI*), start, and stop time (±*SEM*). **(A)** Mean peak time (±*SEM*) from the re-training to the probe session. Open circles denote the aCSF group, and closed squares denote the Pir+AP5 group. An asterisk (*) indicates a significant simple main effect of group [*p* < .050]. **(B)** Mean *DI ±SEM* from the re-training to the probe session. Mean (±*SEM*) start **(C)** and stop **(D)** times at the baseline, shift, and probe sessions.

##### 3.2.2.3. DI

The mean *DI*s (± *SEM*) from the first re-training session to the probe session are shown in Figure 5B. In all sessions, the mean *DI*s of the Pir+AP5 group were slightly lower than those of the aCSF group. A mixed two-way ANOVA in the re-training sessions showed a significant main effect of the group [*F* (1,11) = 8.108, *p* = .016, *ηG^2^* = .107], but not the main effect of the session and the interaction [*F* (2.06,22.66) = 3.106, *p* = .063, *ηG^2^* = .191; *F* (2.06,22.66) = .256, *p* =.783, *ηG^2^* = .019, respectively]. A mixed two-way ANOVA from the baseline to the probe sessions revealed a significant main effect of the session [*F* (1.31,14,44) = 38.4921, *p* < .001, *ηG^2^* = .671], but not the main effect of the group and the interaction [*F* (1,11) = .017, *p* = .900, *ηG^2^* < .001, *F* (1.31,14.44) = 1.628, *p* = .228, *ηG^2^* = .079]. In multiple comparisons of the session, the *DI* of the probe session was significantly higher than that of the baseline and shift sessions [*p_s_* < .009].

##### 3.2.2.4. The number of responses per sec

The mean number of responses per second (± *SEM*) in the baseline, shift, and probe sessions are shown in Table 1. There was no noticeable difference between the groups in any of the sessions. A mixed two-way ANOVA revealed a significant main effect of the session [*F* (2,22) = 14.591, *p* < .001, *ηG^2^* = .367], but not the main effect of the group and the interaction [*F* (1,11) = .071, *p* = .795, *ηG^2^* = .004; *F* (2,22) = .805, *p* = .460, *ηG^2^* = .031, respectively]. In multiple comparisons of the session, the value of the probe session was significantly lower than that of the baseline and shift sessions [*p_s_* < .006].

#### 3.2.3. Trial by trial analysis

The mean start and stop times (± *SEM*) are shown in Figure 5C and D, respectively. Unlike the peak times, there were no noticeable differences between the groups in the baseline, shift, and probe sessions in both indices. A mixed two-way ANOVA of the start times showed a significant main effect of session [*F* (2,22) = 10.449, *p* < .001, *ηG^2^* = .277], but not the main effect of the group and the interaction [*F* (1,11) = .233, *p* = .639, *ηG^2^* = .013, *F* (2,22) = 1.193, *p* = .322, *ηG^2^* = .042, respectively]. In multiple comparisons of the sessions, the start time of the shift and probe sessions were significantly higher than that of the baseline session [*p_s_* < .006]. For the stop times, the main effect of the session was significant [*F* (1.17,12.82) = 47.544, *p* < .001, *ηG^2^* = .740], but not the main effect of the group and the interaction [*F* (1,11) = 1.013, *p* = .336, *ηG^2^* = .031, *F* (1.17,12.82) = 3.191, *p* = .094, *ηG^2^* = .160, respectively]. In multiple comparisons of the sessions, the stop time of the shift and probe sessions was significantly higher than that of the baseline session, and that of the probe session was significantly lower than that of the shift session [*p_s_* < .003].

### 3.3. Discussions

The results of the shift session suggest that the co-inhibition of M1 and NMDA receptors did not impair the acquisition of new-duration memory. The peak times, start time, and stop time of both groups significantly increased from baseline to the shift session (Figure 5A, C, and D). Moreover, the difference in these indices and *DI* between the groups was not significant in the shift session. These results suggest that 1) new duration memory was acquired in both groups, and 2) the strength of memory acquisition in both groups was similar to each other.

Interestingly, the results of the probe session suggest that the co-inhibition of M1 and NMDA receptors impairs the consolidation of new-duration memory. The rats in both groups acquired new duration memory in the shift session. However, the next day, the peak time of the Pir+AP5 group in the probe session was significantly lower than that of the aCSF and Pir+AP5 groups in the shift session (Figure 5A). These findings suggest that the new duration memory of the Pir+AP5 group was poorer than that of the aCSF group. This phenomenon is consistent with the operational definition of impairment of memory consolidation [6]. Unlike the peak times, the differences in the *DI* and *R^2^* between the groups in the probe session were not significant (Figure 5B). Moreover, the difference in the number of responses per second between groups was also not significant in the shift and probe sessions. Therefore, it is suggested that the memory impairment in the Pir+AP5 group cannot be explained by the side effect of the poor precision of interval timing and/or motivational factors. Taken together, the findings of Experiment 2 suggest that the intra-DS co-infusion of pirenzepine and D-AP5 before the shift session prevented the acquired duration memory from being consolidated.

## 4. Experiment 3: Effect of the intra-DS infusion of solo or mixed M1 and NMDA receptor antagonist on the consolidation of duration memory

The results of Experiment 2 suggested that co-inhibition of M1 and NMDA receptors impaired the consolidation of duration memory. However, this effect has not been replicated and could be caused by the inhibition of M1 or NMDA receptors. In Experiment 3, we aimed to confirm the replicability of the findings of Experiment 2 and examine the effect of the sole infusion of the drugs on the consolidation of duration memory. All experimental procedures were approved by the Doshisha Committee of Animal Experiments (A20078).

### 4.1. Material and methods

The materials and methods were the same as in Experiment 2, except for the following points. Another cohort of 48 naïve male Wistar albino rats, aged approximately 11 weeks, was used. Surgery was performed before the training began. In training, the ratio of the food and empty trials was 42:18 until session 20 and 15:15 after session 21. In the shift session, sole or mixed D-AP5 (3.0 μg/side) and/or pirenzepine (10.0 μg/side) were infused other than aCSF.

#### 4.1.1. Dependent variables

In addition to the dependent variables in Experiments 1 and 2, we calculated the percentage of change rates of the peak time in all combinations of the baseline, shift, and probe sessions. Other than Bonferroni’s method, Dunnett’s method was used to make pairwise multiple comparisons between the aCSF group and other groups. In addition, a one-sample t-test was used. In the one-sample t-test, the *p*-values were adjusted by *p* × 6, the number of all comparisons, to avoid the underestimation of type I error.

### 4.2. Results

Eleven rats were removed from the analysis: two died intraoperatively, three were from the removal of cannulas, one from bent injection cannula and bleeding from the guide cannula. Four subjects whose *R^2^* in the probe session was below −2 *SD*, and one whose peak time in the probe session was over ±2 *SD*. As a result, the number of subjects was nine in the aCSF, 11 in the AP5, nine in the Pir, and eight in the Pir+AP5 groups.

#### 4.2.1. Histology

A representative section and all internal cannula tips are shown in Figures 1A and D, respectively. All internal cannulas were inserted into the DS.

#### 4.2.2. Session by session analysis

##### 4.2.2.1. Response rate distributions

The mean response rate distributions and their fitted curves are shown in Figure 6. In the baseline session (Figure 6A, D, G), the distributions overlapped with the peaks at approximately 17 s. In the shift session (Figure 6B, E, H), the distributions shifted rightward compared to the baseline session. The peaks of the aCSF and Pir+AP5 groups were located at approximately 27 s, and those of the AP5 and Pir groups were located at approximately 29 s. In the probe session, the distributions of the AP5 and aCSF groups overlapped as a whole, whereas those of the Pir and Pir+AP5 groups were located leftward compared with those of the aCSF groups. The peak of the aCSF, AP5, Pir, and Pir+AP5 groups in the probe session were approximately 24, 28, 17, and 19 s, respectively. The mean *R^2^* (± *SEM*) values are shown in Table 1. The values were greater than .799 in all sessions and groups. A mixed two-way ANOVA (group × session) showed a significant main effect of session [*F* (2,66) = 12.843, *p* < .001, *ηG^2^* = .154], but not the main effect of the group and the interaction [*F* (3,33) = .500, *p* = .685, *ηG^2^* = .024; *F* (6,66) = .070, *p* = .999, *ηG^2^* = .003]. In multiple comparisons for the session, the *R^2^s* of the shift and probe sessions was significantly lower than that of the baseline session [*p_s_* < .001].

**Figure 6.**
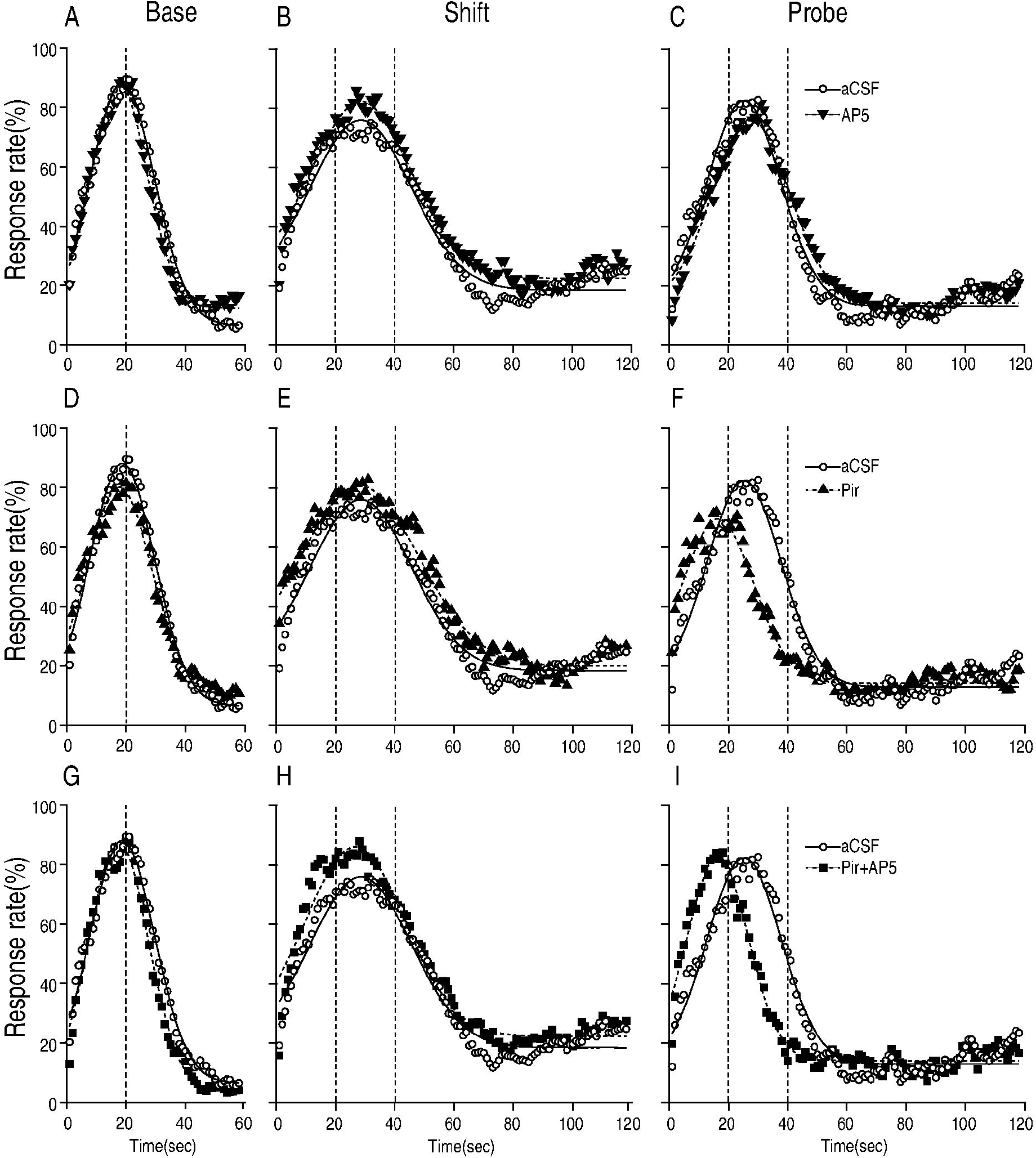
Mean response rate distributions and fitted Gaussian curves. Shown are the aCSF group vs AP5 **(A-C)**, Pir **(D-F)**, and Pir+AP5 **(G-I)**. Open circles denote the aCSF group in all panels. Closed inverted triangles show the AP5 group, closed triangles the Pir group, and closed squares the Pir+AP5 group. Dotted lines show the target duration of the PI-20 s and PI-40 s sessions.

##### 4.2.2.2. Peak time

The mean peak times (± *SEM*) in session 25 to the probe session are shown in Figure 7A. During sessions 25-30, the peak times were consistently approximately 17 s. A mixed two-way ANOVA in sessions 25 to 30 showed neither a significant main effect nor the interaction [for group, *F* (3,33) = .015, *p* = .998, *ηG^2^* = .001; for session, *F* (5,165) = 2.098, *p* = .068, *ηG^2^* = .013; for interaction, *F* (15,165) = 1.171, *p* = .299, *ηG^2^* = .021]. The mean peak times (± *SEM*) in the baseline and shift sessions were indifferent between the groups, whereas those of the Pir and Pir+AP5 groups in the probe session were lower than those in the aCSF group. A two-way mixed ANOVA of group and session (baseline, shift, and probe) showed a significant main effect of session and interaction [*F* (2,66) = 34.592, *p* < .001, *ηG^2^* = .342; *F* (6,66) = 3.377, *p* = .006, *ηG^2^* = .132], but not the main effect of the group [*F* (3,33) = 2.481, *p* = .078, *ηG^2^* = .102]. The simple main effect of the group in the probe session [*F* (3,33) = 9.812, *p* < .001, *ηG^2^* = .472] and of the session in all groups were significant [aCSF, *F* (2,16) = 10.389, *p* = .001, *ηG^2^* = .265; AP5, *F* (2,20) = 16.906, *p* < .001, *ηG^2^* = .448; Pir, *F* (2,16) = 9.087, *p* = .002, *ηG^2^* = .460; Pir+AP5, *F* (2,14) = 10.140, *p* = .002, *ηG^2^* = .470]. In multiple comparisons of the groups, the peak times of the Pir and the Pir+AP5 groups in the probe session were significantly lower than those in the aCSF group [Dunnett’s method, *p*_s_ < .017]. In multiple comparisons of the session in the aCSF and AP5 groups, the peak times of the shift and probe sessions were significantly higher than those in the baseline session [aCSF, *p_s_* < .029; AP5, *p_s_* < 0.003]. In multiple comparisons of the sessions in the Pir and Pir+AP5 groups, the peak time of the shift session was significantly higher than that of the baseline and probe sessions [Pir, *p_s_*< .033; Pir+AP5, *p_s_*< .016].

**Figure 7.**
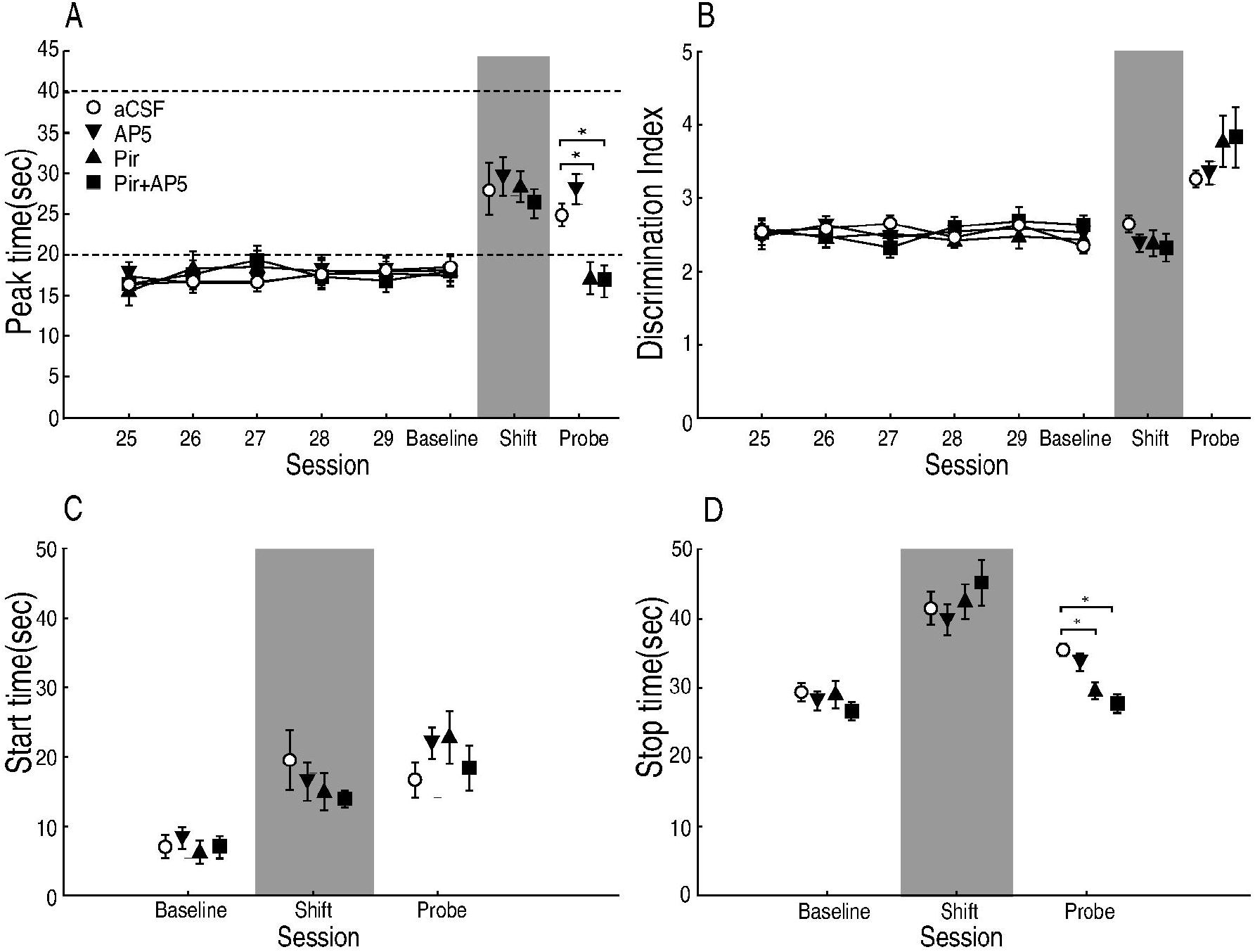
Mean peak time, discrimination index (*DI*), start, and stop time (±*SEM*). **(A)** Mean peak time (±*SEM*) from session 25 to the probe session. Open circles denote the aCSF group in all panels. Closed inverted triangles denote the AP5 group, closed triangles the Pir group, and closed squares the Pir+AP5 group. An asterisk (*) indicates a significant difference *vs*. the aCSF group [Dunnett’s method, *p* < .050]. **(B)** Mean *DI* ±*SEM* from the re-training to the probe session. Mean (±*SEM*) start **(C)** and stop **(D)** times at the baseline, shift, and probe sessions.

##### 4.2.2.3. DI

The mean *DI*s (± *SEM*) in session 25 to the probe session are shown in Figure 7B. Although the mean *DI*s varied slightly, there was no difference between the groups. A two-way mixed ANOVA in sessions 25 to 30 showed neither significant main effects nor interaction [for group, *F* (3,33) = 0.138, *p* = .937, *ηG^2^* = .007; for session, *F* (5,165) = .710, *p* = .617, *ηG^2^* = .010; for interaction, *F* (15, 165) = 1.207, *p* = .271, *ηG^2^* = .048]. The mean *DI*s of all groups in the shift and probe sessions were shifted in the same way. A two-way mixed ANOVA of the group and session (baseline, shift, and probe) showed a significant main effect of session [*F* (1.42, 46.73) = 75.935, *p* < .001, *ηG^2^* = .507], but not the main effect of the group and the interaction [*F* (3, 33) = .627, *p* = .603, *ηG^2^* = .031. *F* (4.25, 46.73) = 2.360, *p* = .064, *ηG^2^* = .087]. In multiple comparisons, the *DI* of the probe session was significantly higher than that of the baseline and shift sessions [*p_s_* < .001].

##### 4.2.2.4. The number of responses per sec

The mean number of responses per second (± *SEM*) in the baseline, shift, and probe sessions are shown in Table 1. The values in the shift and probe sessions tended to be lower than those in the baseline session. A mixed two-way ANOVA showed a significant main effect of session [*F* (2,66) = 87.553, *p* < .001, *ηG^2^* = .307], but not the main effect of the group and the interaction [*F* (3,33) = .909, *p* = .447, *ηG^2^* = .064, *F* (6,66) = .661, *p* = .682, *ηG^2^* = .010, respectively]. In multiple comparisons of the sessions, all differences in all combinations were significant [*p_s_* < .002].

##### 4.2.2.5. Change rate

The mean percentage of change rates (± *SEM*) of the shift/baseline, probe/shift, and probe/baseline are shown in Figure 8. For the shift/baseline, the values were indifferent irrespective of the session and drugs used. A two-way ANOVA (with or without Pir × with or without AP5) showed neither a significant main effect nor the interaction [for the effect of Pir, *F* (1, 33) = .009, *p* = .925, *ηG^2^* < .001; for AP5, *F* (1,33) = .007, *p* = .935, *ηG^2^* < .001; interaction, *F* (1,33) = 1.061, *p* = .311, *ηG^2^* = .031]. One-sample t-tests of the Pir+ (combination of the Pir and Pir+AP5 groups) and Pir-(combination of the aCSF and AP5 groups) groups showed that the values were significantly higher than 100% [one-tailed, Pir+: *t* (19) = 6.551, *p* < .001; Pir-: *t* (16) = 4.869, *p* < 001]. For the probe/shift, the change rates of the Pir+ group were lower than those of the Pir+ group. A two-way ANOVA showed a significant main effect of Pir [*F* (1,33) = 14.054, *p* < .001, *ηG^2^* = .299], but not the main effect of AP5 and the interaction [*F* (1,33) = .097, *p* = .757, *ηG^2^* = .003; *F* (1,33) = .030, *p* = .864, *ηG^2^* < .001, respectively]. One-sample t-tests showed that the value was significantly lower than 100% in the Pir+ group [*t* (19) = −.640, *p* > 1], but not in the Pir-group [*t* (16) = −5.227, *p* < .001]. For the probe/baseline, the change rates of the Pir+ group were lower than those of the Pir-group. A two-way ANOVA showed a significant main effect of Pir [*F* (1,33) = 13.223, *p* < .001, *ηG^2^* = .286], but not the main effect of AP5 and the interaction [*F* (1, 33) = .622, *p* = .436, *ηG^2^* = .019; *F* (1, 33) = .949, *p* = .337, *ηG^2^* = .028, respectively]. One-sample t-tests showed that the value was significantly higher than 100% in the Pir-group [*t* (19) = 6.063, *p* < .001], but not in the Pir+ group [*t* (16) = .122, *p* > 1].

**Figure 8.**
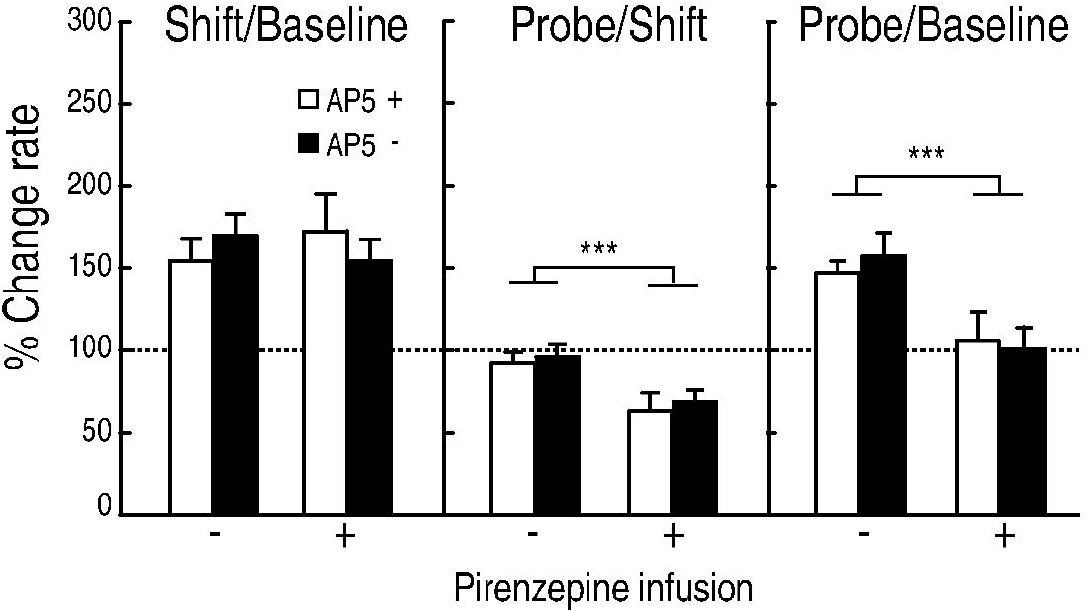
Mean change rate (±*SEM*) of the shift/baseline, probe/shift, and probe/baseline. White and black bars denote the presence (+) or absence (-) of D-AP5, respectively. Plus and minus on the X-axis label denote the presence or absence of pirenzepine, respectively. The horizontal dotted line indicates 100%. Asterisks (***) indicate a significant main effect of Pir [*p* < .001].

#### 4.2.3. Trial by trial analysis

The start and stop times (± *SEM*) are shown in Figures 7C and D. Although the start times of all sessions were indifferent between all groups, the stop times of the Pir and Pir+AP5 groups in the probe session were lower than those in the aCSF group. A mixed two-way ANOVA showed a significant main effect of the session and the interaction [*F* (2, 66) = 50.469, *p* < .001, *ηG^2^* = .338; *F* (6, 66) = 1.732, *p* = .127, *ηG^2^* = .050, respectively], but not the main effect of the group [*F* (3,33) = .220, *p* = .892, *ηG^2^* =.013]. In multiple comparisons of the sessions, the start time of the shift and probe sessions was higher than that of the baseline session [*p_s_* < .001]. For the stop times, the main effect of the session and interaction was significant [*F* (2,66) = 113.758, *p* < .001, *ηG^2^* = .598; *F* (6, 66) = 3.710, *p* = .003, *ηG^2^* = .127, respectively], but not the main effect of the group [*F* (3, 33) = .478, *p* = .699, *ηG^2^* = .024]. The simple main effect of the group in the probe session [*F* (3, 33) = 8.484, *p* < .001, *ηG^2^* = .435], and of the session in all groups were significant [aCSF: *F* (2,16) = 19.915, *p* < .001, *ηG^2^* = .539; AP5: *F* (2, 20) = 19.259, *p* < .001, *ηG^2^* = .474; Pir: *F* (2,16) = 36.100, *p* < .001, *ηG^2^* = .613; Pir+AP5: *F* (2, 14) = 42.670, *p* < .001, *ηG^2^* = .732]. In multiple comparisons, the stop times of the Pir and Pir+AP5 groups in the probe session were significantly lower than those in the aCSF group [Dunnett’s method, *p_s_* < .017]. The stop time of the aCSF group in the shift session was significantly higher than that of the baseline and probe sessions [*p*_s_ < .020]. The stop time of the AP5 group in the shift session was significantly higher than that in the baseline and probe sessions [*p*_s_ < .015]. The stop times of the Pir and Pir+AP5 groups in the shift session were significantly higher than those in the baseline and probe sessions [Pir, *p_s_* < .002; Pir+AP5, *p_s_* < .001].

### 4.3. Discussion

The results of the shift session suggested that the sole or co-inhibition of M1 and/or NMDA receptors did not impair the acquisition of new-duration memory. The peak times and start/stop times of all groups significantly increased from baseline to the shift session (Figure 7A, C, D, and 8). Moreover, there was no significant difference between groups in these indices, the *DI*s, and the number of responses per second (Figure 7B and Table 1). These results suggest that 1) new duration memory was acquired in all groups, and 2) the strength of the memory acquired was similar to each other.

The results of the probe session suggested that if only M1 receptors were inhibited, the consolidation of new-duration memory was impaired. In the probe session, the peak and stop times of the Pir and Pir+AP5 groups, but not the AP5 group, were lower than those of the aCSF group (Figure 7A and D). Therefore, the results of Experiment 2 were replicated in the Pir+AP5 group. Moreover, the ANOVA of change rates revealed a significant main effect of pirenzepine, but not AP5 treatment (Figure 8). Additionally, the change rate of the pirenzepine-treated groups was significantly lower than 100% in the probe/shift comparison, whereas the value was not significantly higher than 100% in the probe/baseline comparison.

This means that the sole inhibition of M1 receptors was sufficient to impair the consolidation of duration memory. The difference in the *DI* and the number of responses per second between the groups were also not significant in all sessions. Therefore, it is suggested that the impairment of memory consolidation in the Pir and Pir+AP5 groups cannot be explained by the side effects of the poor precision of interval timing and/or motivational factors.

## 5. General discussion

The aim of the present study was to investigate the effect of inhibition of D1, M1, and NMDA receptors in the DS on the consolidation of duration memory. In Experiment 1, the co-inhibition of D1 and NMDA receptors had no effect on either the acquisition or consolidation of duration memory. In Experiment 2, the co-inhibition of M1 and NMDA receptors impaired the consolidation, but not the acquisition, of duration memory. In Experiment 3, the effect of co-inhibition of M1 and NMDA receptors was replicated. More importantly, the sole inhibition of M1, but not NMDA receptors, impaired the consolidation of duration memory. These results strongly suggest that M1 receptors in the DS are required for consolidation of duration memory in the range of interval timing.

The impairment of consolidation of duration memory by the inhibition of M1 receptors might be caused by the suppression of LTP or the occurrence of LTD in MSNs. Ninety-five percent of DS neurons consist of MSNs [33]. Moreover, M1 receptors have been identified in three-quarters of MSNs [34]. MSNs are innervated by cholinergic interneurons within the DS and cholinergic neurons of the pedunculopontine (PPT) and laterodorsal tegmental nuclei (LDT) [35]. A physiological study suggested that the inactivation of M1 receptors of MSNs reduced the cortico-striatal glutamatergic synaptic transmission and promoted LTD [12, 36]. Conversely, other studies have shown that the inhibition of M1 receptors inhibits LTP [18], and their activation by cholinergic interneurons is required for the long-term excitation of MSNs [37]. These studies suggest that the impairment of the consolidation of duration memory was caused by promoted LTD or inhibited LTP in MSNs with the inactivation of M1 receptors.

The impairment of behavioral flexibility by the inhibition of M1 receptors cannot explain our findings. Our procedure required rats to replace the initial duration memory with a new memory under the inhibition of M1 receptors. Previous studies have shown that the inhibition of dorsomedial striatal M1 receptors in rats impairs reversal learning, but not the initial acquisition [38,39]. The procedure of the previous studies is similar to ours in that the initial memory is replaced with another memory under M1 receptor blockade. Therefore, it can be hypothesized that the inhibition of M1 receptors impairs reversal learning, not the consolidation of memory. However, we disagree with this explanation. In previous studies [38, 39], M1 blockers were infused before a reversal learning session, and the effect was exerted in the session. In contrast, the M1 blocker was infused in the shift session (corresponding to the reversal learning session of the previous studies), but the effect was not exerted in the session in our experiment. Therefore, we believe that the “reversal learning hypothesis” can be rejected.

From a theoretical point of view, the ineffectiveness of dopamine and glutamate receptor blockers is surprising but not unreasonable. The SBF model proposed that MSNs become coincidence detectors of cortical glutamatergic inputs with the coactivation of dopaminergic synapses from the midbrain. Although D1 and NMDA receptors are one of the key players underlying the neuronal plasticity in MSNs [12-17; for review, see 40], other types of receptors such as dopamine D2 and/or metabotropic glutamate receptors are also involved in the plastic changes of MSN synapses [40]. Therefore, our findings do not contradict the prediction of the SBF model and narrow the candidates for the receptor types inducing neuronal plasticity underlying the formation of duration memory.

## 6. Conclusion

In conclusion, our results suggest that M1, but not D1 and NMDA, receptors in the DS are important for the consolidation of duration memory.

## Author contributions

M. N.: software, formal analysis, investigation, writing of the original draft, writing of the review, and editing. T.K.: supervision and investigation. T.K., A.N., N.S., and A.K.: investigation. T.H.: conceptualization, methodology, software, investigation, writing the review and editing, supervision, and project administration.

## Acknowledgement

We would like to thank Editage (www.editage.com) for English language editing.

## Data availability statement

The dataset and log of analysis are available from the Open Science Framework (OSF) (https://osf.io/yrkg3/?view_only=adbb1a3d994f4b04bc5f85a82d22eaa4).

## Abbrevations

SBF: Striatal Beat Frequency
DS: dorsal striatum
MSNs: medium spiny neurons
HFS: high frequency stimulus
ERK: extracellular signal-regulated kinase
Arc: activity-regulated cytoskeletal-associated protein kinase
PI: peak interval
DI: discrimination index
Pir: pirenzepine
PPT: pedunculopontine
LDT: laterodorsal tegmental nuclei

